# Droplet-compatible single-cell DNA methylation sequencing with PreTIC

**DOI:** 10.64898/2026.04.15.718726

**Authors:** Sheng Zhang, Faming Wang, Yan Zhang, Keng-Jung Lee, Carl Engman, Hong-Zhang He, Yong Fan, Si-Yang Zheng

**Affiliations:** Department of Biomedical Engineering, Carnegie Mellon University, Pittsburgh, PA, USA; Allegheny Health Network Cancer Institute, Pittsburgh, PA, USA; Captis Diagnostics, Inc, Pittsburgh, PA, USA; Department of Electrical and Computer Engineering, Carnegie Mellon University, Pittsburgh, PA, USA

## Abstract

Single-cell DNA methylation sequencing has remained technically specialized due to challenges of interfacing conversion chemistry with droplet microfluidics. We introduce pre-tagmentation in situ conversion (PreTIC), which enables whole-methylome profiling on a commercial droplet platform using off-the-shelf reagents. With PreTIC, we produce over 13,000 single-cell methylomes from fixed frozen cells in two days and resolve cell-specific epigenetic variation in human peripheral blood mononuclear cells.

## Main text

DNA methylation is a central regulator of gene expression and cellular identity^1^, and single-cell DNA methylation sequencing (scDNAm-seq) enables the analysis of epigenetic heterogeneity that is obscured in bulk assays^2^. Despite its importance, scDNAm-seq remains less accessible than single-cell RNA-seq or ATAC-seq. Existing approaches rely primarily on plate-based workflows with limited throughput^3,4^ or combinatorial indexing strategies^5–7^ requiring custom reagents, labor-intensive protocols and extended experimental timelines. Droplet-based scDNAm-seq^8^ has progressed more slowly due to the technical complexity of methylation conversion chemistries, which involve DNA denaturation and multiple buffer exchanges.

In contrast, droplet platforms like 10x have widely enabled scRNA-seq and scATAC-seq through standardized systems. Extensions of 10x scATAC-seq now support high-throughput single-cell whole-genome sequencing^9^, as well as chromatin conformation capture^10^, underscoring the versatility of droplet-based technologies mediated by in situ tagmentation, a process that targets intact nuclei and double-stranded DNA. However, bisulfite conversion (or enzymatic conversion^11^) requires single-stranded DNA substrates and generates single-stranded products. This mismatch represents a major obstacle to adapting existing scATAC-seq methods for scDNAm-seq.

Here we introduce <u>pre</u>-tagmentation in situ <u>c</u>onversion (PreTIC), a strategy that establishes a workflow fully compatible with in situ tagmentation and droplet barcoding, and performed entirely in bulk (Fig. 1a). In PreTIC, genomic DNA undergoes denaturation and bisulfite conversion inside individual nuclei, followed by in situ second-strand synthesis to regenerate double-stranded DNA suitable for standard tagmentation. To preserve nuclear integrity under the harsh chemical conditions of bisulfite conversion, we implement hydrogel embedding adapted from expansion microscopy^12^. PreTIC produces intact nuclei containing converted double-stranded DNA and allows standard in situ tagmentation directly. As a proof of concept, we deploy PreTIC on the commercial 10x ATAC platform to enable scDNAm-seq without custom oligonucleotides or dedicated microfluidics. The entire procedure takes about two days from fixed cells or nuclei to final sequencing libraries (Supplementary Figs. 1-3). It enables profiling of over 10,000 cells per sample without cell multiplexing, and the throughput can be readily scaled up by utilizing additional channels.

**Figure 1.**
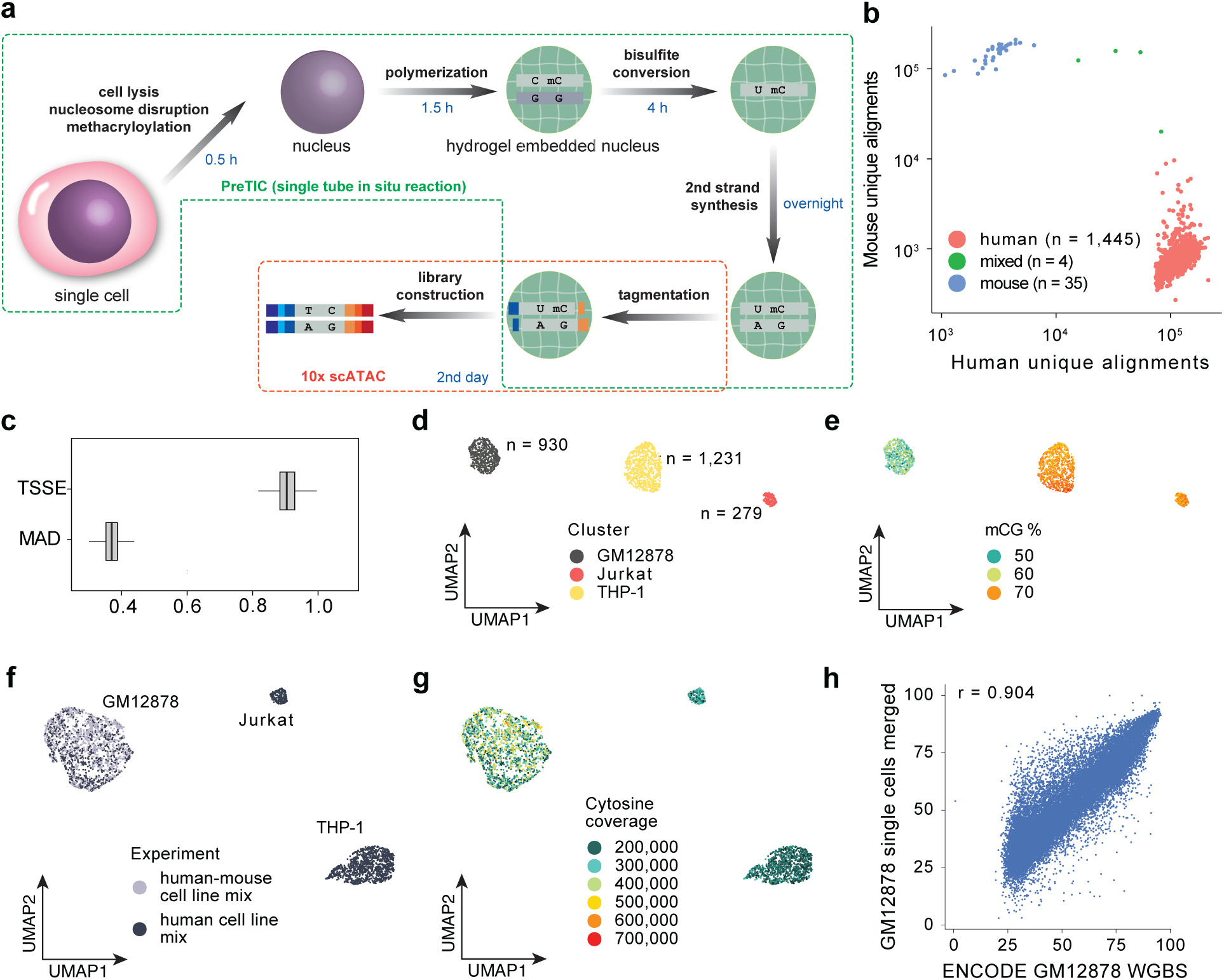
| Overview and validation of scDNAm-seq with PreTIC. **a**, Schematic of scDNAm-seq with PreTIC. Genomic DNA inside fixed, hydrogel-embedded nuclei undergoes denaturation and in situ bisulfite conversion, followed by second-strand synthesis to restore double-stranded DNA prior to tagmentation. The converted nuclei are then processed using the standard 10x Genomics ATAC workflow to generate single-cell DNA methylation libraries. **b**, Scatterplot of single-cell libraries showing counts of unique human and mouse alignments, for estimation of barcode collision. **c**, Boxplots of transcription start site enrichment (TSSE) and median absolute deviation (MAD) score per cell for GM12878, for assessment of coverage uniformity. For all boxplots, center lines denote medians; boxes indicate the 25th and 75th percentiles; whiskers extend to 1.5× the interquartile range. **d,e**, UMAP projection of the human cell line mix colored by cell type assignment (d) or global CG methylation (mCG) level (e). **f,g**, Integrated UMAP of human cells from the human-mouse mixture and the human cell line mixture, colored by experimental run (f) or cytosine coverage (g). **h**, Correlation between ENCODE bulk GM12878 WGBS data and merged GM12878 single-cell methylomes (n windows = 25,929).

We first conducted a species-mixing experiment to investigate the purity of scDNAm profiles. From a human-mouse mixture, we recovered 1,484 single-cell methylomes (mean 134,540 unique alignments per cell), of which 1,445 mapped predominantly to human GM12878 cells and 35 to mouse A20 cells, with 4 mixed-species barcodes corresponding to an estimated doublet rate of 5.84% (Fig. 1b). GM12878 cells exhibited uniform genome-wide coverage without transcription start site enrichment and displayed median absolute deviation scores comparable to single-cell whole-genome sequencing studies (Fig. 1c and Supplementary Fig. 4), supporting even genomic representation^9,13^.

We next profiled a mixture of three human cell lines (GM12878 B cells, Jurkat T cells, and THP-1 monocytes) to evaluate the method’s ability to discriminate between cell types. A total of 2,440 cells were recovered, and UMAP projection revealed clear separation of linages (Fig. 1d,e), even at a low sequencing depth (mean 96,820 unique reads). GM12878 cells from independent experiments co-clustered despite differences in sequencing depth (Fig. 1f,g), demonstrating reproducibility. Pseudobulk CG methylation (mCG) profiles of the GM12878 single-cell cluster showed strong correlation with ENCODE whole-genome bisulfite sequencing (r = 0.904; Fig. 1h).

We further characterized the performance of the assay on fixed frozen peripheral blood mononuclear cells (PBMCs). Libraries were generated from maximal cell input loaded into a single 10x Genomics Chromium channel and sequenced to a depth of 4.2 billion read pairs. After adapter trimming, 99.63% reads were retained, and 86.40% aligned with minimal strand bias (50.49% aligned to Watson strand) (Fig. 2a,b). 13,701 single-cell methylomes were recovered with a mean of 353,998 unique reads per cell, capturing 1,582,307 cytosines and 84,136 CG sites (Fig. 2c, Supplementary Fig. 5). Coverage scaled proportionally with unique read counts (Supplementary Fig. 6). Read alignment rate showed no correlation with CG coverage or mCG levels, and mitochondrial contamination was minimal (Supplementary Figs. 7,8). The global CH methylation (mCH) level was 6.04% (Fig. 2d), modestly elevated relative to expectations from mammalian somatic cells^14^. This residual cytosine non-conversion is consistent with the challenge of maintaining complete DNA denaturation during in situ bisulfite treatment. Five clusters were identified, corresponding to major PBMC populations: CD8^+^ T, CD4^+^ T, natural killer (NK), monocyte (Mono) and B cells (Fig. 2e). Global mCG levels varied across clusters (Supplementary Fig. 9a), with monocytes and NK cells skewed toward increased methylation and lymphoid populations displaying broader distributions. This shift persisted at megabase-scale resolution, while overall methylation architecture remained largely conserved (Supplementary Fig. 10). In contrast, global mCH rates, cytosine coverage and alignment rates were similar across clusters (Supplementary Fig. 9b–d), suggesting that these technical metrics do not account for the observed separation. Cluster-aggregated mCG profiles across canonical lineage genes revealed locus-level patterns consistent with expected immune identities (Fig. 2f and Supplementary Fig. 11). Differential promoter mCG at established marker genes further distinguished PBMC subtypes (Fig. 2g). Previously reported PBMC differentially methylated regions (DMRs) were recapitulated, providing independent validation of cell-type assignments (Fig. 2h)^15^. In total, 117,047 cluster-specific DMRs (adjusted p ≤ 0.05) were identified, 78.11% of which overlapped candidate cis-regulatory elements annotated by SCREEN (Fig. 2i). Gene set enrichment analysis of these regions highlighted immune-related processes, including B cell activation and T cell differentiation (Fig. 2j), supporting their functional relevance.

**Figure 2.**
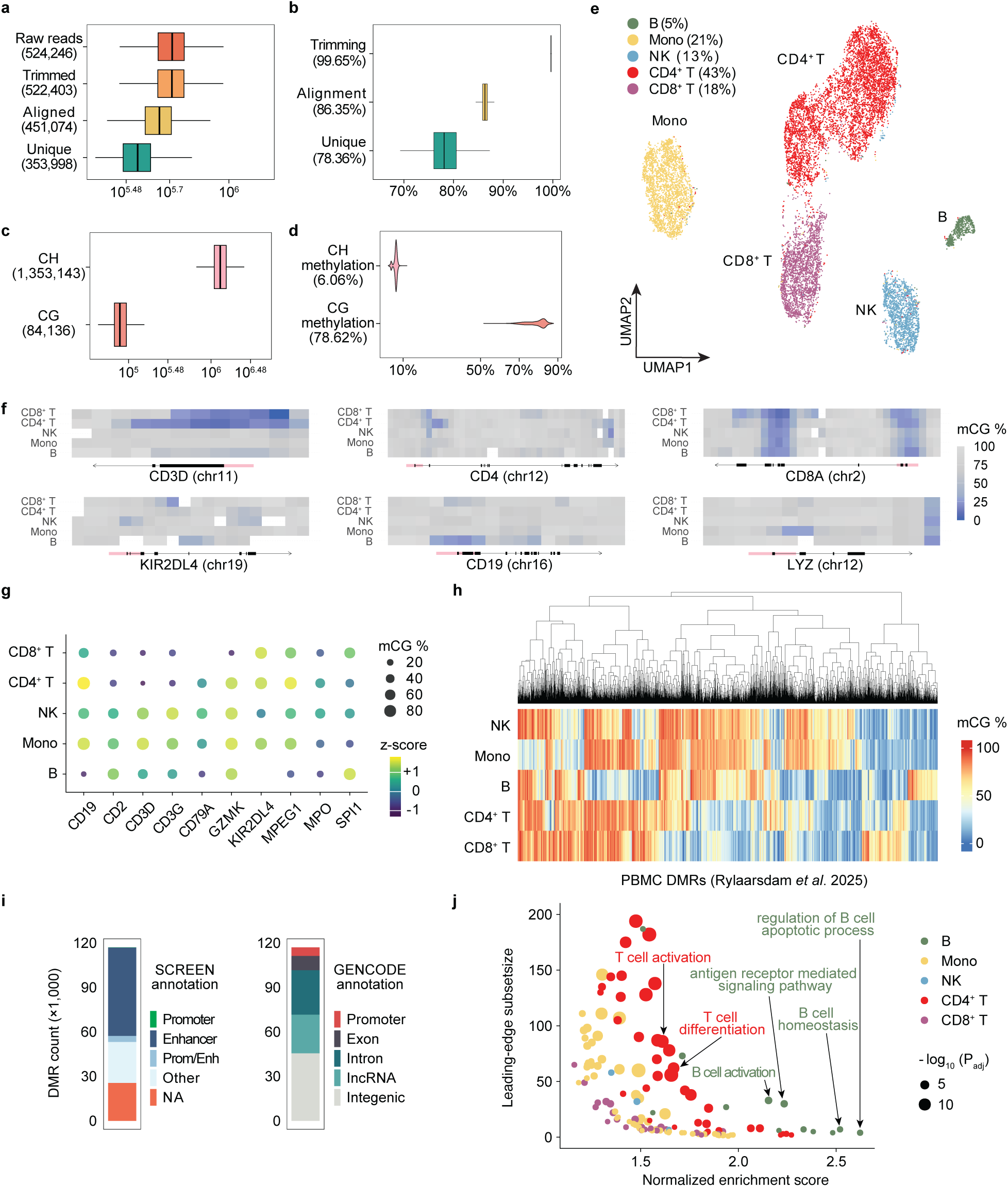
| Performance of scDNAm-seq with PreTIC on human peripheral blood mononuclear cells. **a,b**, Number and percentage of reads retained per cell after each preprocessing step; mean values are shown in parentheses. **c,d**, Cytosine coverage and methylation rates in CH and CG contexts. **e**, UMAP projection of PBMCs colored by cluster identity. **f**, Heatmaps showing CG methylation levels at 1-kb resolution across selected canonical marker loci distinguishing pan-T cells, CD4⁺ T cells, CD8⁺ T cells, NK cells, B cells and monocytes in cluster-aggregated PBMC profiles. **g**, Promoter CG methylation levels of PBMC marker genes across aggregated cell clusters. **h**, Heatmap of CG methylation rates across aggregated cell clusters for the top 3,000 most variable DMRs previously described by Rylaarsdam *et al.* 2025. **i**, Distribution of DMRs stratified by SCREEN and GENCODE annotations. **j**, Gene set enrichment analysis using cluster-specific DMRs.

PreTIC is optimized for high-throughput single-cell methylome profiling at moderate sequencing depth. By reconciling bisulfite conversion with in situ tagmentation, the method operates on standardized commercial droplet infrastructure without custom oligonucleotide design or dedicated microfluidic systems. Using a single 10x Genomics Chromium channel, PreTIC recovers over 10,000 single-cell methylomes with high alignment rates and uniform genome-wide representation. Relative to previously described droplet-based whole-methylome methods^8^, this approach achieves improved alignment efficiency and genomic coverage while simplifying experimental implementation (Supplementary Table 1).

## Methods

### Human blood collection

This study was approved by the Carnegie Mellon University Institutional Review Board (IRB ID STUDY2020_00000127) and all participants provided informed consent. Peripheral blood from a healthy male donor was collected in EDTA-coated tubes and processed within one hour of collection.

### Cell culture

Cell lines (GM12878, Coriell; A20, ATCC TIB-208; Jurkat, Clone E6-1, ATCC TIB-152; THP-1, ATCC TIB-202) were maintained at 37 °C in a humidified incubator containing 5% CO₂. GM12878 cells were grown in RPMI 1640 (Cytiva SH30027.02) supplemented with 15% (v/v) FBS (VWR 97068-085), 1× penicillin-streptomycin-glutamine (Gibco 15140122). Jurkat and THP-1 cells were grown in RPMI 1640 (Cytiva SH30027.02), supplemented with 10% (v/v) FBS, and 1× penicillin-streptomycin-glutamine. A20 were grown in RPMI 1640 (ATCC 30-2001), supplemented with 10% (v/v) FBS, and 1× penicillin-streptomycin-glutamine. Cultured cells were harvested and washed with PBS (Corning 21-040-CV) before proceeding to fixation.

### Human peripheral blood mononuclear cell (PBMC) isolation

PBMCs were isolated from whole blood using SepMate™ tubes (STEMCELL Technologies 85450) and Lymphoprep™ (STEMCELL Technologies 18061) according to the manufacturer’s instructions. Briefly, blood was diluted 1:1 with room temperature PBS and layered directly into SepMate tubes pre-filled with Lymphoprep. Samples were centrifuged at 1200 × g for 10 minutes at room temperature with the brake on. The PBMC layer was poured off into a new tube, washed twice with PBS, and centrifuged at 300 × g for 8 minutes at room temperature for all wash steps. Cell counts and viability were assessed using trypan blue (Sigma-Aldrich T8154) exclusion and a hemocytometer.

### Single-cell library preparation

Single cells in suspension were fixed in 0.8% PFA/PBS (Santa Cruz SC281692) for 15 minutes at room temperature. Fixed cells were pelleted, resuspended in freezing buffer (20 mM HEPES-NaOH pH 8.0, 10 mM NaCl, 10% (v/v) DMSO) and stored at -80°C in aliquots with slow freezing. All centrifugation steps were performed at 300-500 × g for 5-10 minutes at room temperature. On the day of the experiment, the cells were thawed from -80 °C and pelleted. Approximately two millions of cells were resuspended in 200 µL LA buffer (20 mM HEPES-NaOH pH 8.0, 10 mM NaCl, 0.05% (w/v) sodium dodecyl sulfate (SDS, Fisher BP2436200), 12.5 mM methacrylic acid N-hydroxysuccinimide ester (MA-NHS, Sigma-Aldrich 730300)) and incubated for 30 minutes at room temperature for concurrent cell lysis (nuclei release), nucleosome disruption (to open up the chromatin) and methacryloylation. In situ polymerization was followed by adding a 100 µL mixture of monomers and initiators to a final concentration of 1% (w/v) acrylamide (AAm, RPI A11260), 1% (v/v) N,N-dimethylacrylamide (DMAA, Sigma-Aldrich 274135), 0.3% (w/v) ammonium persulfate (APS, Sigma-Aldrich A3678) and 0.2% (v/v) tetramethylethylenediamine (TEMED, Bio-Rad 1610800). After polymerization under vacuum for 1.5 hours at room temperature, the reaction was mixed with 1 mL TT (10 mM Tris-HCl pH 8.0, 0.1% (v/v) Tween 20 (Sigma-Aldrich P2287)) and centrifuged. The supernatant was carefully removed, leaving a residual volume of 30-50 µL to avoid disturbing the nuclei pellet. Formamide (Roche 11814320001) was added to achieve a final concentration of 85% (v/v). Nuclei were then resuspended and heated for 20 minutes at 85 °C to denature DNA. 900 µL SB solution (2.5 M sodium metabisulfite (Thermo Fisher 419582500), adjusted to pH ∼5.0 with 10 M NaOH, 20 mM hydroquinone (Thermo Fisher A11411.30), 8% glycerol (Fisher BP229-4)) was added and the reaction was transferred to 65 °C and incubated for 2 hours. 160 µL ethanol (Decon 2701) was then added to induce precipitation, which lowers viscosity of the solution and facilitates pelleting of bisulfite treated nuclei. After centrifugation, the pellet was resuspended in 1 mL 0.35 M NaOH supplemented with 0.1% (v/v) Triton X-100 (Sigma-Aldrich T8787) and incubated for 20 minutes at room temperature for desulfonation. The nuclei were washed three times with 1 mL TTB (10 mM Tris-HCl pH 8.0, 0.1% (v/v) Tween 20, 0.04% (w/v) bovine serum albumin (BSA, Sigma-Aldrich A7030)). Washed nuclei were counted. 200,000 to 400,000 nuclei were taken and distributed to PCR tubes with 25,000 nuclei in each well containing 1.25 µL 10 mM dNTP mixture (Bio Basic DD0056) and 4 µl 50 ng/µL random hexamers (IDT 51-01-18-01). The tubes were incubated on ice for 5 minutes before heated at 65 °C for 5 minutes and placed on ice for at least two minutes. A master mix containing rCutSmart™ Buffer (NEB B6004) and Klenow Fragment (3’→5’ exo-) (NEB M0212M), was then added and the reaction (final 25 µL per well containing 0.5 mM dNTP, 8 ng/µL random hexamers, 25,000 cells, 1 U/µL Klenow Fragment (3’→5’ exo-), 1× rCutSmart buffer) was incubated at 4 °C for 15 minutes, and 37 °C (ramp 1 °C/s) overnight (16-20 hours) in a thermocycler. The reactions were afterwards pooled, diluted by 1 mL TTB, and centrifuged. The pelleted nuclei were washed with 1 mL TTB, filtered through a Flowmi™ Cell Strainer (40 microns, Bel-Art H136800040), and resuspended in 1× Diluted Nuclei Buffer (10x Genomics PN-2000207) before submitted to Single Cell Core at the University of Pittsburgh for 10x Genomics Single Cell ATACv2 (10x Genomics PN-1000390). 10x libraries were prepared following the manufacturer’s instructions, except that the Barcoding Enzyme (PN 2000125/2000139) supplied in the kit was substituted with Q5U® Hot Start High-Fidelity DNA Polymerase (NEB M0515) to tolerate uracil residues.

### Library quantification and sequencing

Libraries were quantified with Qubit dsDNA HS Assay Kit (Thermo Fisher Q32854). The fragment size was determined using Agilent High Sensitivity D5000 ScreenTape® System (Agilent 5067-5592) (Supplementary Fig. 3). Libraries were sequenced at Novogene Corporation Inc. on an Illumina NovaSeq X Plus using 25B or 10B flow cells, with paired-end reads (150 cycles for Read 1, 8 cycles for Index 1, 16 cycles for Index 2, and 150 cycles for Read 2).

### Sequencing data processing

FASTQ reads were barcode demultiplexed using unidex (https://github.com/adeylab/unidex) and then trimmed with Trim Galore (v0.6.10) to remove adapter sequences. Trimmed read pairs with at least one read shorter than 20 bp were excluded.

Alignment to the human genome (GRCh38) was performed using BSBolt (v1.6.0) with default parameters. The resulting BAM file was split by barcode. Individual BAM files were coordinate-sorted and deduplicated using SAMtools (v1.22). Unique read counts were used to distinguish single cells from background noise. A bimodal distribution was fitted using *GaussianMixture* from sklearn.mixture (v1.6.1), and a threshold was defined based on the 95% confidence interval of the normal distribution with the higher mean (Supplementary Fig. 5). Barcodes with unique aligned reads below this cutoff were excluded from downstream analyses. Methylation calls were generated from alignments with MAPQ ≥ 10 using premethyst *extract* (https://github.com/adeylab/premethyst).

### Methylation analysis

For the cell line libraries, CG methylation scores were computed over 100 kb non-overlapping windows using amethyst^15^ *makeWindows* (v1.0.4; https://github.com/lrylaarsdam/amethyst). For the PBMC libraries, shrunken means of CG methylation residuals proposed by Kremer et al.^16^ were computed over 100 kb non-overlapping windows using MethSCAN (v1.1.0). Missing values in the resulting feature matrix were replaced with 0, and dimensionality reduction was performed using amethyst *runIrlba*. UMAP projections were generated with the R package uwot (v0.2.4), and Louvain clustering was carried out using the R package igraph (v2.1.4). Single-cell coverage within clusters was aggregated to produce pseudobulk profiles using amethyst *calcSmoothedWindows*. Heatmaps of CG methylation levels across canonical marker genes (Supplementary Fig. 11) were produced with amethyst *heatMap*, and the cell type of each cluster was inferred by visual examination of the methylation patterns. Differentially methylated regions (DMRs) specific to each cluster, defined by comparison against all other clusters, were detected using MethSCAN. DMRs with adjusted p-values ≤ 0.05 were annotated for genomic features using SCREEN^17^ (V4) annotations and GENCODE^18^ (v43) annotations. Gene set enrichment analysis (GSEA) was performed using the fgsea R package (v1.32.2). Gene Ontology Biological Process (GOBP) gene sets were obtained from the MSigDB database via the msigdbr R package (v25.1.1). DMRs overlapping genes were ranked by p-value, and p-values were adjusted for multiple testing using the Benjamini-Hochberg false discovery rate (FDR) procedure. GOBP terms with adjusted p-values ≤ 0.05 were reported.

### Human-mouse mix library development

PFA fixed human (GM12878) and mouse (A20) cell lines were thawed from -80 °C and mixed in an approximate ratio of 4:1, and proceeded with library preparation as described above, except that prior to the final TTB wash and 10x submission, the cells were resuspended in TTB supplemented with 1.25 mM EvaGreen® Dye (Biotium 31000), and subjected to flow sorting. Flow sorting was performed with a BD FACS™ flow sorter in purity mode to collect all single nuclei into a 1.7 mL Eppendorf tube.

In the alignment step, reads were mapped to a human-mouse (GRCh38-GRCm39) hybrid genome, and deduplicated using premethyst *bam-rmdup*. The number of unique alignments to each species was calculated using scitools *barnyard-compare* (https://github.com/adeylab/scitools). Putative single-cell libraries with fewer than 90% of alignments mapping to a single species were classified as detectable inter-species collision events. The observed number of pure and mixed events was used to estimate the total doublet rate under the assumption of random cell pairing. The expected proportion of mixed-species doublets was calculated as 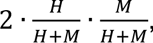, where *H* and *M* are the numbers of observed human and mouse cells, respectively. The total number of doublets was then estimated by dividing the number of observed mixed events by this probability, and the overall collision rate was calculated as the ratio of estimated total doublets to the total number of recovered cells. We note that the human-mouse ratio was dramatically altered in the recovered single cells. Possible reasons include: (1) Different cell cultures may exhibit varying sensitivities to PFA fixation and SDS treatment, which is well documented in the Hi-C literature^19,20^ and has also been observed by us. (2) Flow cytometry gating might introduce bias toward the majority cell population, a limitation that has been noted in previous publications as well^5^. For reanalysis of human GM12878 cells, the FASTQ files were split by species using sciMETv2 *sciMET_speciesSplit.pl* (https://github.com/adeylab/sciMETv2) and FASTQ reads designated to human were re-processed as described above.

### Human cell line mix library development

Three human cell lines were selected: GM12878 (B lymphocyte, female), THP-1 (monocyte, male) and Jurkat (T lymphoblast, male). Fixed cell lines were thawed from -80 °C and mixed in a roughly equal ratio and proceeded with library preparation as described above. Flow sorting was performed as described in human-mouse mix library development. For dimensionality reduction and clustering, the sex chromosome windows were excluded.

### MAD scores

Median Absolute Deviation (MAD) scores were calculated as previously described^9,21^. The autosomal genome was divided into non-overlapping bins of 500 kb. Bin mappability was derived from Bismap^22^ (https://bismap.hoffmanlab.org). Bins with a mappability score < 0.9 or GC content < 20% were excluded. MAD scores for individual cells were then calculated using the following formula^9^:

*MAD score_i_* = *median*(|*d* – *median*(*d*)|), where 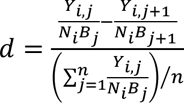, and *Y_i,j_* is the raw read count for the *i*th cell of the *j*th bin, *N_i_* is the cell-specific scaling factor (total reads), and *B_j_* is the bin-specific scaling factor (total reads in bin across all cells).

### TSSE scores

Transcription start site enrichment (TSSE) scores were calculated as previously described^9,21^, by computing the ratio of reads within a 500 bp window centered on annotated transcription start sites (TSSs) to the reads within two 250 bp flanking windows positioned ±1 kb from the TSSs.

### Coverage bias across annotations

ATAC-seq peaks, DNase I hypersensitive sites (DHS), and histone modification ChIP-seq peaks (H3K27ac, H3K27me3, H3K36me3, H3K4me1, H3K4me3, and H3K9me3) for GM12878 cells were downloaded from the ENCODE database. CG island (CGI) annotation was downloaded from the UCSC database. A, B compartment and subcompartment annotation was downloaded from the GEO database^23^. Bedtools (v2.31.0) multicov was used to determine the coverage for each cell across all sites of each annotation bed file. The fraction of total reads per kilobase pair were calculated by summing the coverage and normalizing by the total coverage per cell and the sum of the feature sizes^5^ (Supplementary Fig. 4).

### Correlation with bulk WGBS data

All cells within the GM12878 cluster from the human-mouse mix and cell line mix were collapsed to form a pseudobulk sample. Epigenome Roadmap bulk WGBS data of GM12878 were downloaded from the ENCODE database. Pearson correlation of CG methylation rates across 100 kb non-overlapping windows with ≥ 30 CG measurements were performed.

## Acknowledgements

S.Z. was supported by a scholarship from the Croucher Foundation. Portions of this research were conducted with the advanced computing resources provided by Texas A&M High Performance Research Computing.

