## Supplementary material for "Droplet-compatible single-cell DNA methylation sequencing with PreTIC": supp

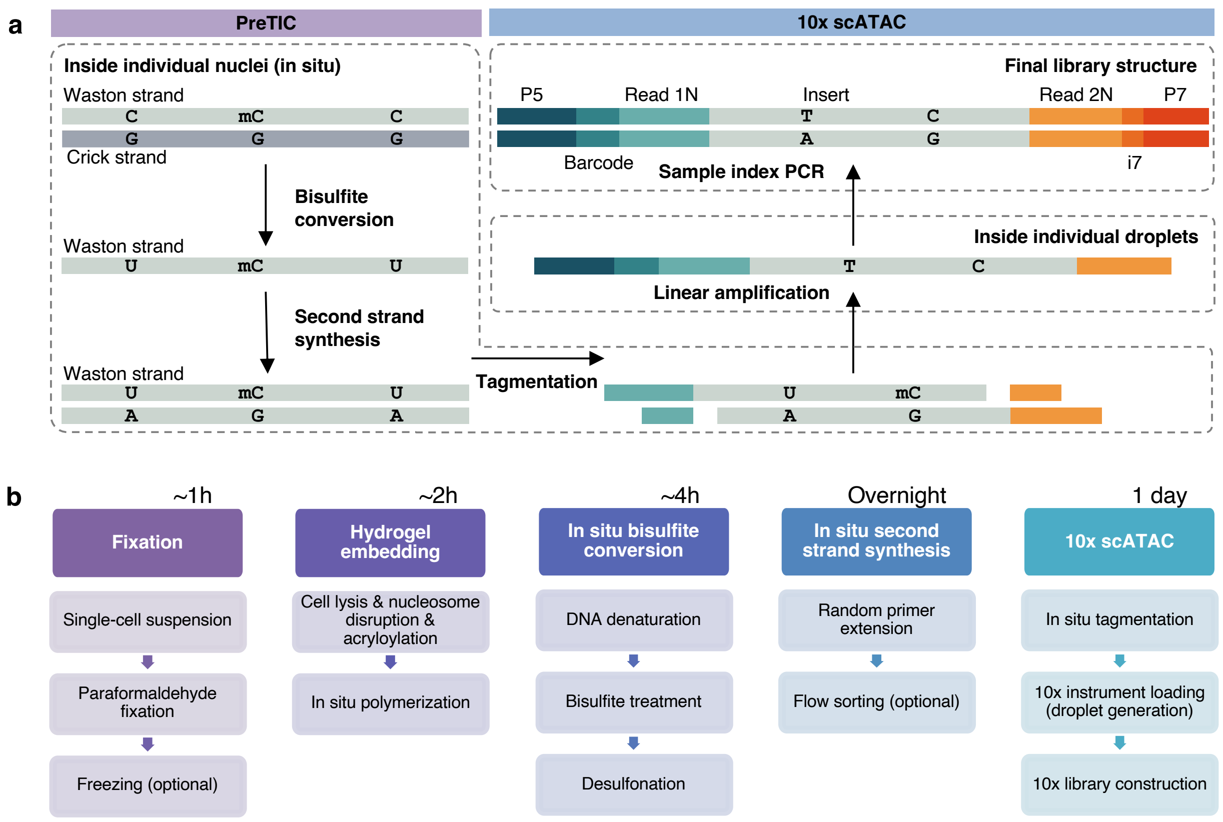


**Supplementary Figure 1** | (a) scDNAm-PreTIC molecular workflow. For simplicity, only Waston strand is shown in full. (b) scDNAm-PreTIC experimental workflow with time estimates.

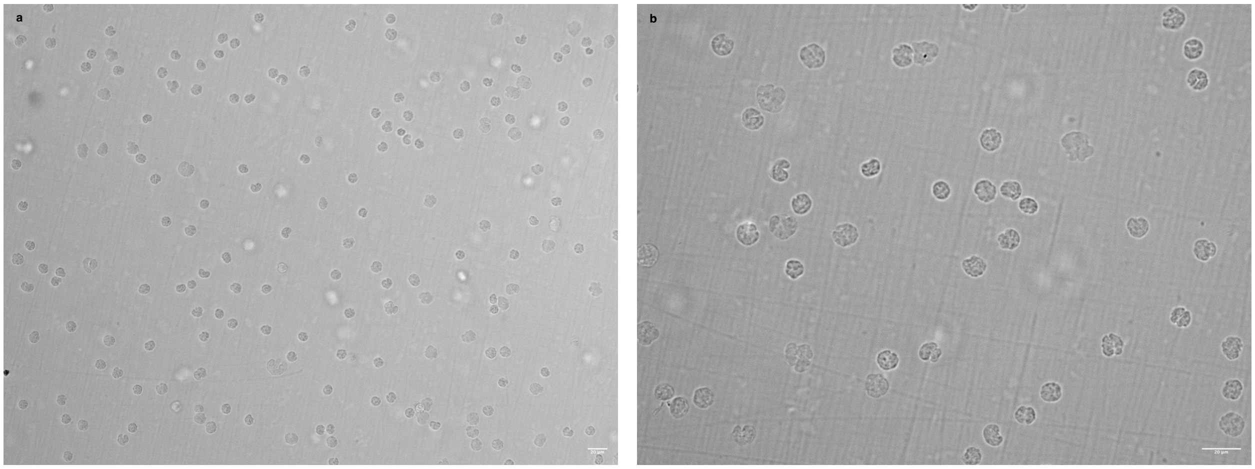


**Supplementary Figure 2** | Brightfield images of PreTIC treated PBMC nuclei, after hydrogel embedding, in situ bisulfite conversion and in situ second-strand synthesis, before submission to 10x ATAC, acquired using (a) a 20× objective and (b) a 40× objective. Imaging was performed on an EVOS M7000 microscope.

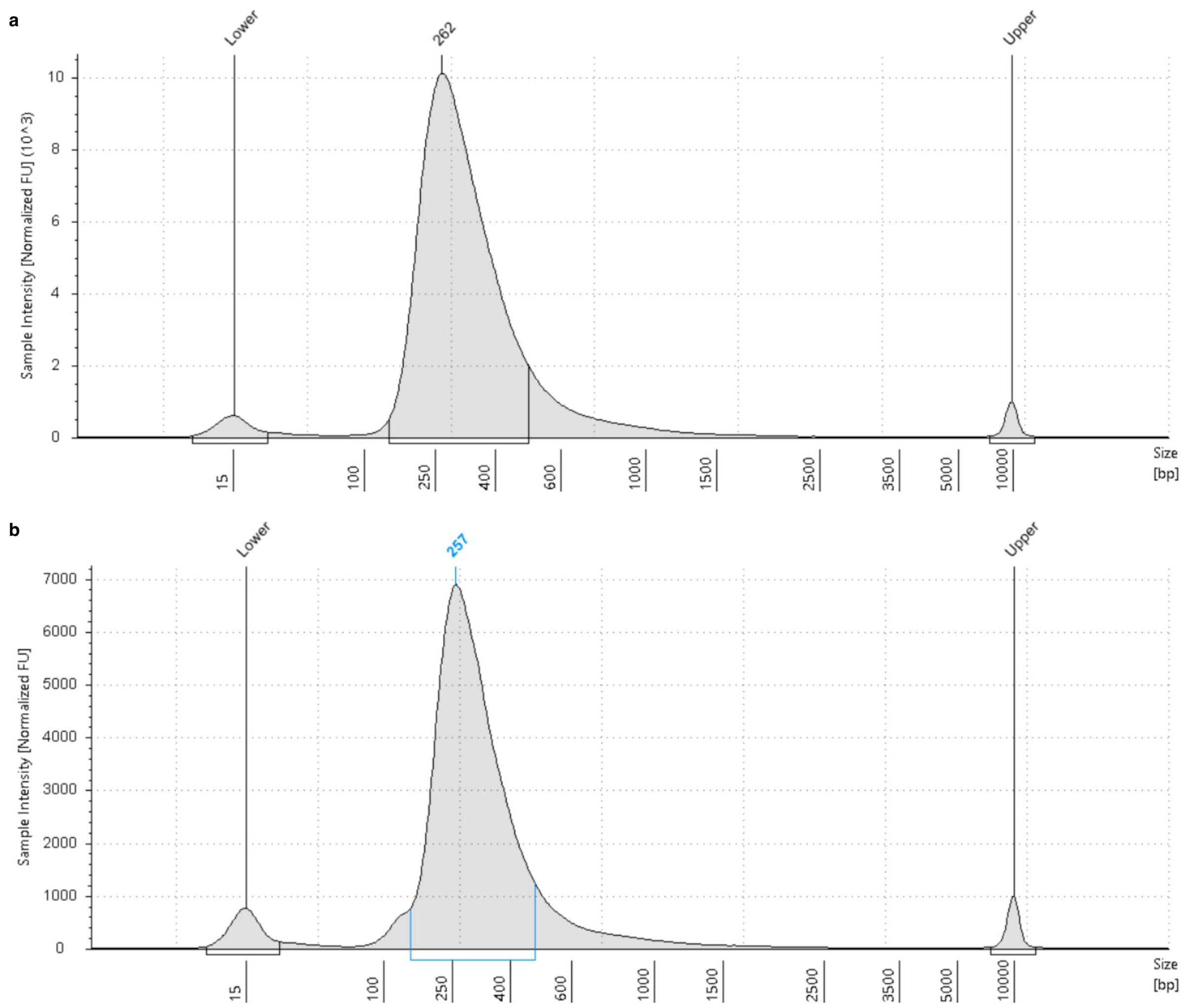


**Supplementary Figure 3** I TapeStation traces of (a) the human cell line mix library and (b) the PBMC library.

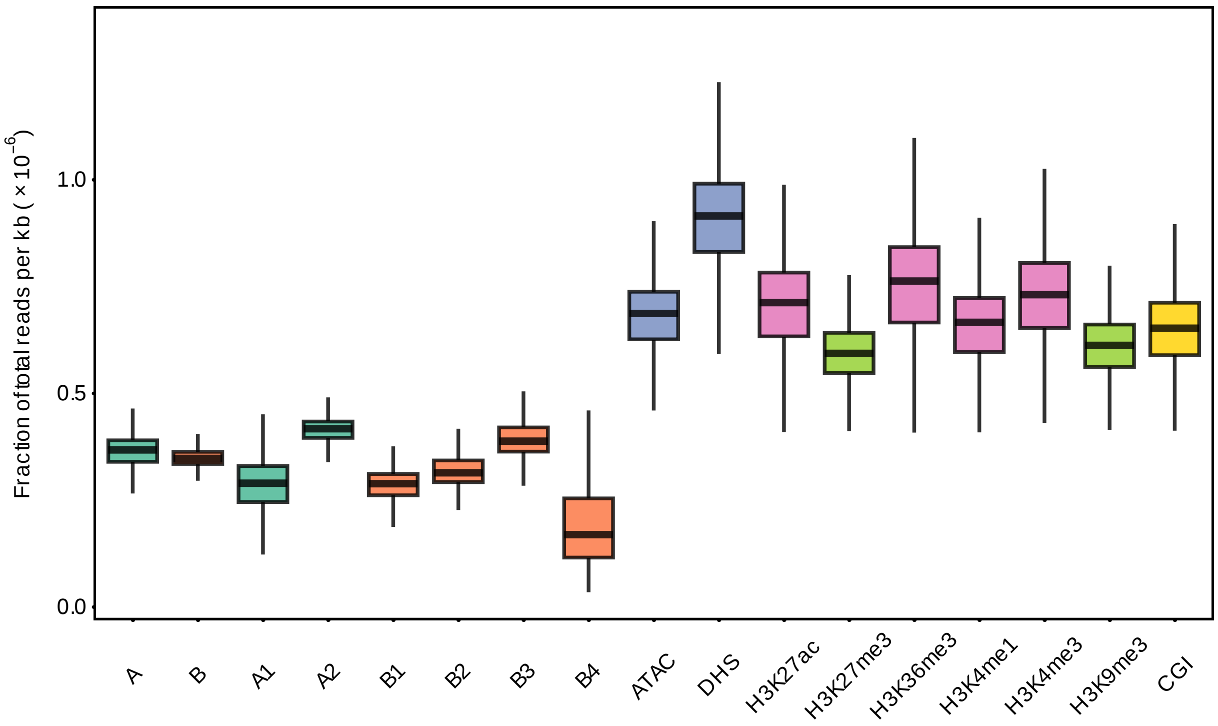


**Supplementary Figure 4** | Coverage of GM12878 single-cell libraries from the human-mouse mix experiment, quantified as the fraction of total reads normalized to feature length (per kilobase), across annotated genomic regions of the GM12878 cell line. A, B: A compartments and B compartments; A1, A2: A subcompartments; B1, B2, B3, B4: B subcompartments; described by Rao *et al.* 2014. ATAC, ATAC-seq peaks. DHS, DNase I hypersensitive sites. CGI, CG island. Slight bias toward open / transcriptional active chromatin was observed.

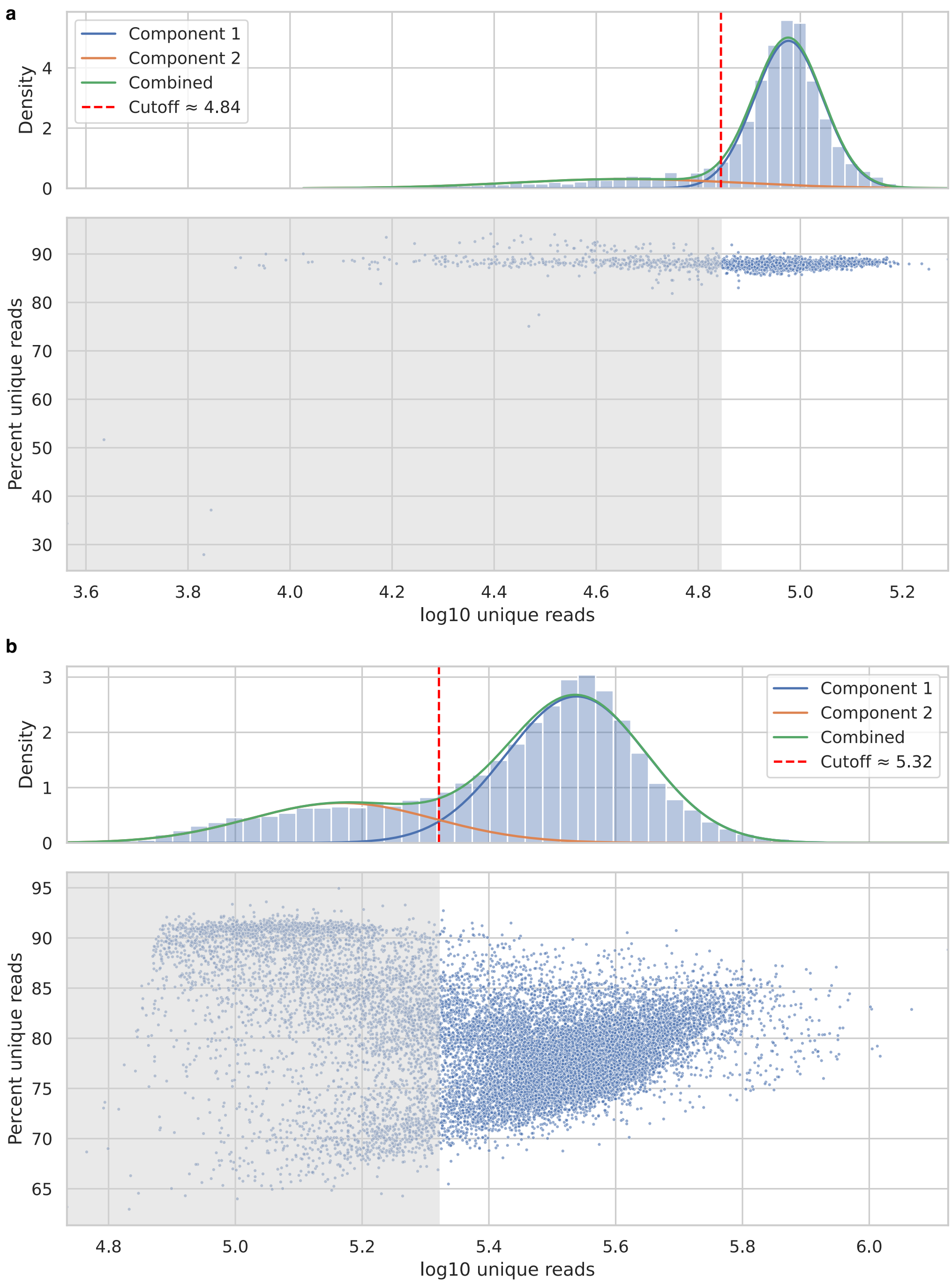


**Supplementary Figure 5** I Library complexity, measured by unique read counts and the percentage of unique reads, and single-cell discrimination based on unique read counts for (a) the human cell line mix experiment and (b) the PBMC experiment. Each dot represents a barcode. Barcodes falling below the unique read count cutoff (grey shaded area) were considered background noise and excluded from downstream analyses. In the human cell line mix experiment, flow cytometry was performed, which likely accounts for the reduced background noise observed.

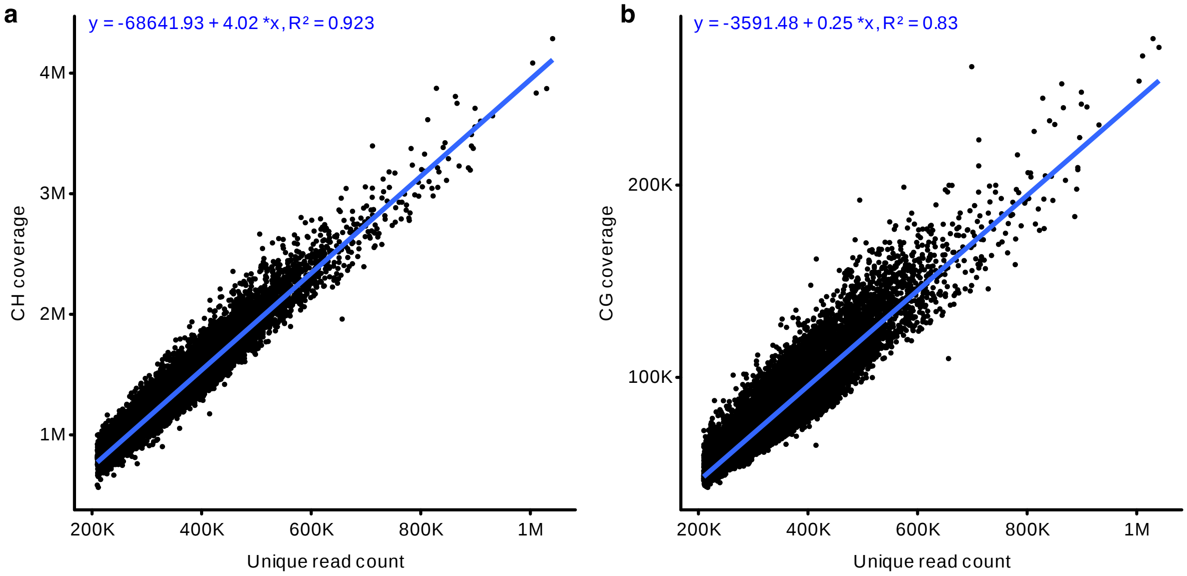


**Supplementary Figure 6** | High correlation between the number of uniquely aligned reads per cell and (a) the number of CH sites or (b) CG sites covered in the PBMC single-cell libraries.

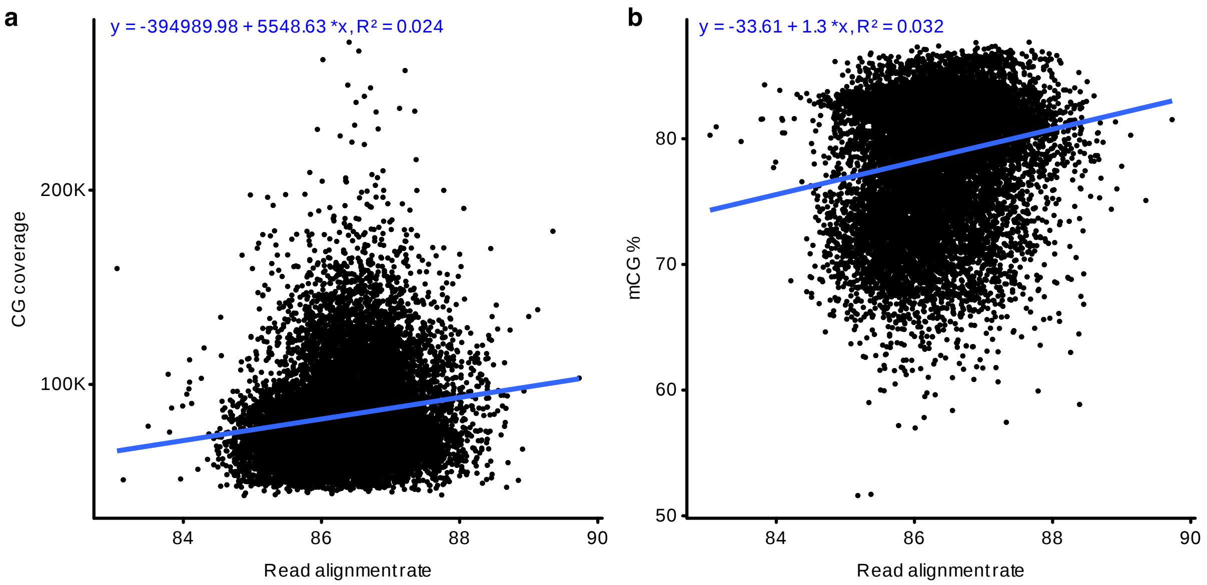


**Supplementary Figure 7** | Low correlation between read alignment rate and (a) CG coverage or (b) CG methylation percentage in the PBMC single-cell libraries.

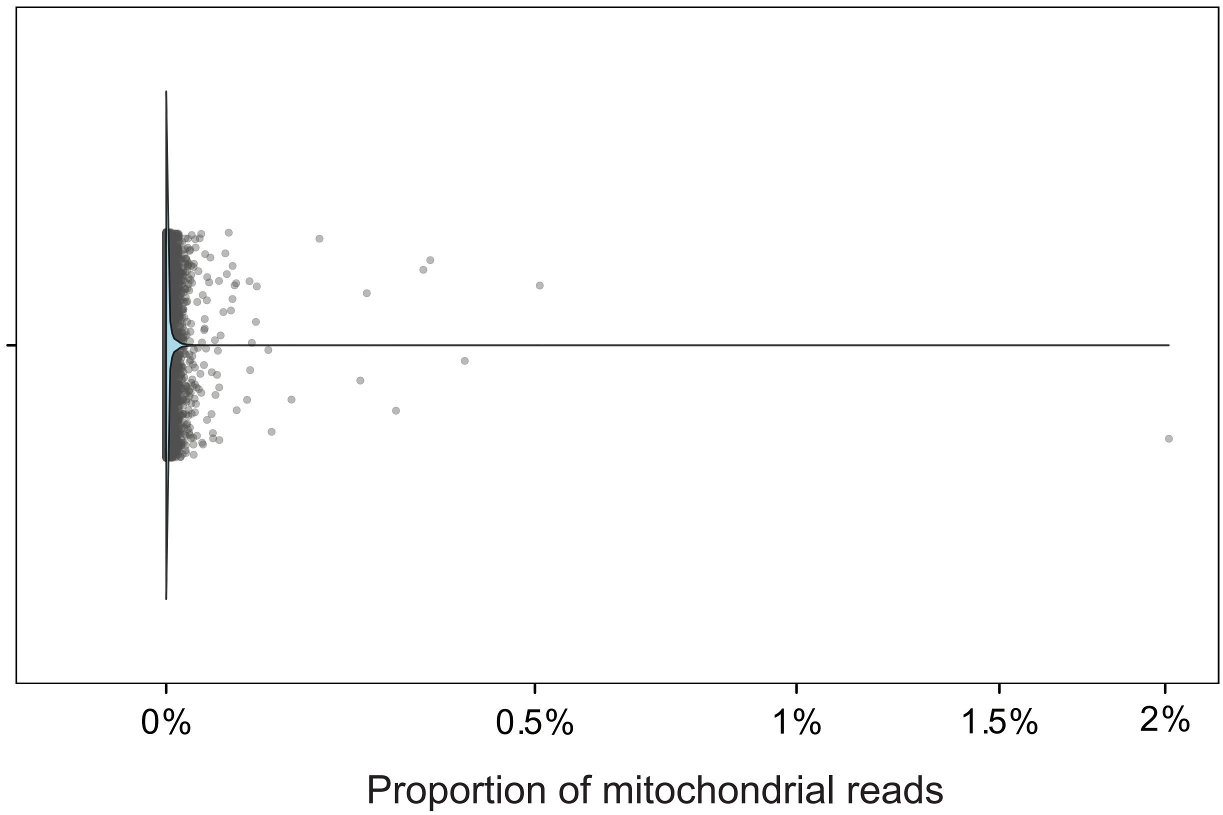


**Supplementary Figure 8** | Low mitochondrial contamination, as measured by the percentage of mitochondrial reads in PBMC single-cell libraries (mean 0.0019%). 7,763 out of 13,701 (56.66%) of cells have no mitochondrial reads.

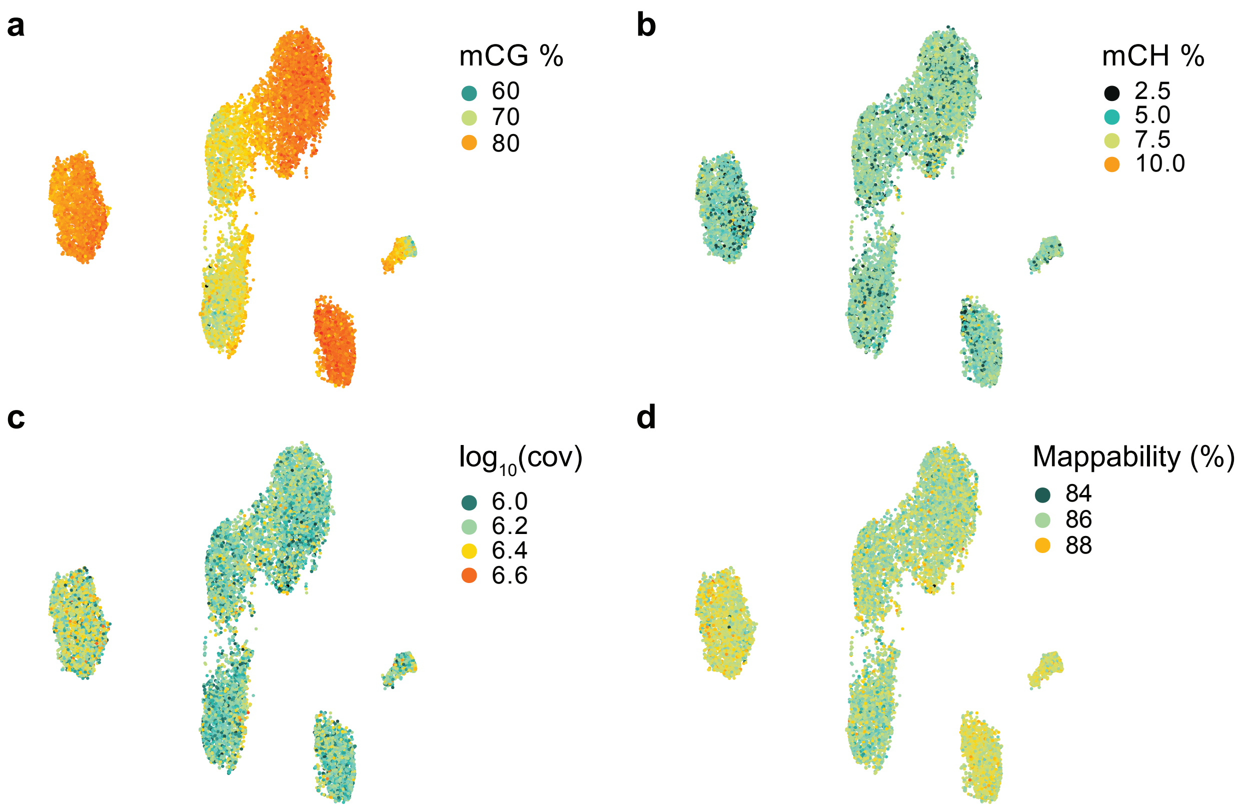


**Supplementary Figure 9** | UMAP projections of the PMBCs colored by (a) global CG methylation level, (b) global CH methylation level, (c) cytosine coverage and (d) read alignment rate.

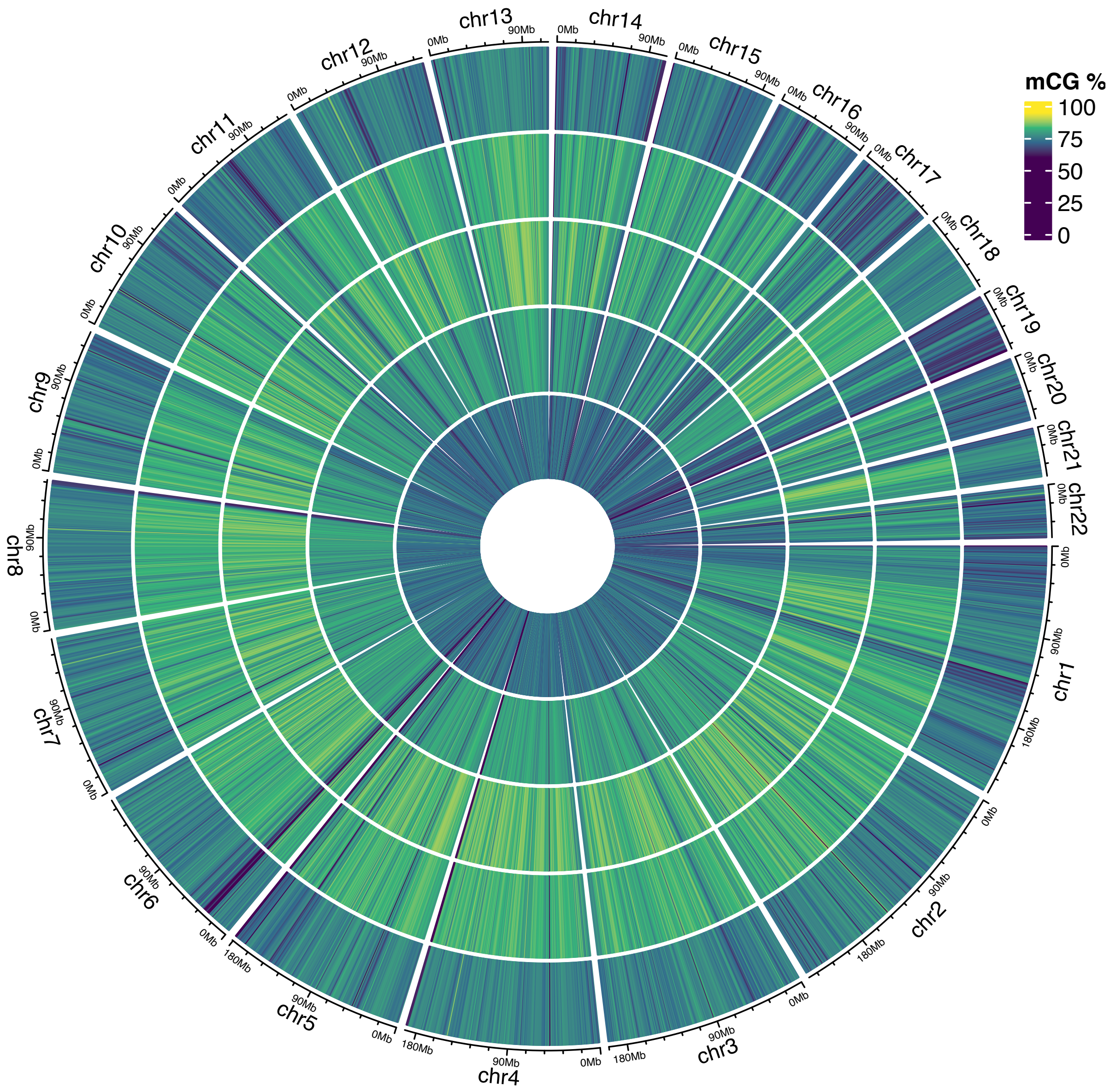


**Supplementary Figure 10** | Circular representation of CG methylation levels across 1-Mb genomic windows in cluster-aggregated PBMC profiles. Tracks are arranged from outermost to innermost as follows: B cells (B), monocytes (Mono), natural killer cells (NK), $CD4^+$ T cells ($CD4^+$T), and $CD8^+$ T cells ($CD8^+$ T).

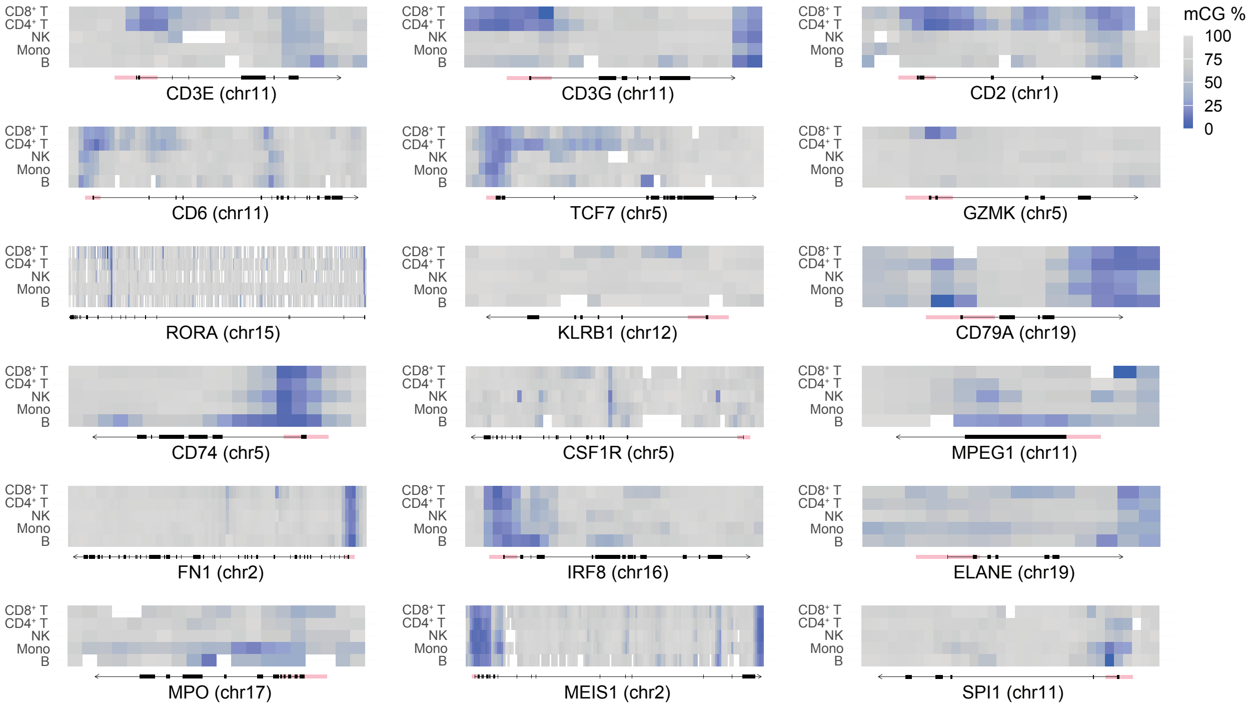


**Supplementary Figure 11** | Heatmaps of CG methylation levels across canonical marker genes in cluster-aggregated PBMC profiles, calculated in 1-kb windows.

| Method | PreTIC (this study) | Drop-BS | sciMETv2 | sciMETv3 |
|---|---|---|---|---|
| **Format** | Droplet (10x ATAC) | Custom droplet microfluidics | Combinatorial indexing | Combinatorial indexing |
| **Cells per run** | >10,000 per 10x channel | ~10,000 | ~1,000 | >10,000 |
| **Mean unique reads per cell (mean raw reads per cell)** | ~$3.5\times10^5$ (~$5.2\times10^5$) | ~$4.1\times10^4$ (~$2.3\times10^5$) | ~$3.1\times10^6$ (~$7.5\times10^6$) | ~$2.6\times10^5$ (~$3.2\times10^5$) |
| **Mean CGs per cell** | ~$8.4\times10^4$ | ~$2.4\times10^4$ | ~$2.2\times10^6$ | NA |
| **Mean cytosines per cell** | ~$1.6\times10^6$ | NA | ~$4.5\times10^7$ | $1.8\times10^6$ |
| **Alignment rate** | 86% | 65% | 69% | 94% |
| **Custom oligos** | No | Yes | Yes | Yes |
| **Custom microfluidic system** | No | Yes | No | No |
| **Compatible with commercial droplet platform** | Yes | No | No | No |

**Supplementary Table 1** | Comparison of high-throughput single-cell DNA methylation sequencing methods. NA, not available.
